# Transcription factor fluctuations underlie cell-to-cell variability in a signaling pathway response

**DOI:** 10.1101/2022.11.30.518555

**Authors:** Avinash Ramu, Barak Cohen

## Abstract

Stochastic differences among clonal cells can initiate cell fate decisions in development or cause cell-to-cell differences in the responses to drugs or extracellular ligands. We hypothesize that some of this phenotypic variability is caused by stochastic fluctuations in the activities of transcription factors. We tested this hypothesis in NIH3T3-CG cells using the response to Hedgehog signaling as a model cellular response. Here we present evidence for the existence of distinct fast and slow responding substates of NIH3T3-CG cells. These two substates have distinct expression profiles, and fluctuations in the activity of the *Prrx1* transcription factor (TF) underlie some of the differences in expression and responsiveness between fast and slow cells. We speculate that similar variability in other TFs may underlie other phenotypic differences among genetically identical cells.

## Introduction

Stochastic molecular fluctuations can cause phenotypic variability among genetically identical cells in the same environment. For example, cell-to-cell fluctuations of SCA1 help determine the lineage along which hematopoietic stem cells will differentiate (Chang et al. 2008), and cell-to-cell fluctuations of EGFR and AXL underlie the resistance of rare melanoma cells to chemotherapy (Shaffer et al. 2017, 2018). Because such molecular fluctuations can have important phenotypic consequences, a key question that needs to be answered is what are the molecular mechanisms underlying cell-to-cell variability within clonal cell populations?

Several mechanisms underlying cell-to-cell variability in gene expression have been identified to date. Some variables that can cause differences between genetically identical cells include cell-cycle stage(Zopf et al. 2013), cellular volume(Kempe et al. 2015; Padovan-Merhar et al. 2015), mitochondrial content(Neves et al. 2010), ribosome numbers (Guido et al. 2007) and cell state(Topolewski et al. 2022; Kiviet et al. 2014; Iwamoto, Shindo, and Takahashi 2016). However, there are likely additional mechanisms controlling cell-to-cell variability in gene expression, and the advent of single-cell genomics(Trapnell 2015; Eling, Morgan, and Marioni 2019) may provide approaches for identifying these mechanisms.

Transcription factors control gene expression and play a vital role in development. Stochastic fluctuations in the activities of TFs can be caused by fluctuations in their expression, localization, or post-translational modification. We hypothesize that stochastic fluctuations in the activities of TFs, and the subsequent fluctuations of their downstream target genes, contribute to cell-to-cell differences among genetically identical cells. A key prediction of this hypothesis is that the expression variability of certain TFs, or that of their target genes, will correlate with phenotypic differences between clonal cells. Cells that stochastically express high amounts of a TF’s target genes might behave differently from clonal cells expressing lower amounts of the same genes.

We use the cellular response to extracellular Hedgehog (HH) as a phenotype to test these predictions. HH signaling polarizes developing tissues through a signaling pathway that results in the activation of the Gli family of TFs (J. H. Kong, Siebold, and Rohatgi 2019; Briscoe and Thérond 2013; Lee, Zhao, and Ingham 2016). HH signaling can therefore be measured using cells carrying a genome-integrated Gli-responsive reporter gene (Pusapati et al. 2018). However, we found that even among clonal cells, treatment with HH results in significant cell-to-cell variation in the activation of a Gli-responsive reporter gene. We therefore attempted to identify TFs whose fluctuations might account for these cell-to-cell differences in HH responsiveness using single-cell RNA sequencing (scRNA-seq). We identified fast- and slow-responding subsets of cells characterized by distinct expression profiles that were caused, in part, by stochastic fluctuations in the activity of the *Prrx1* TF. We found that over-expression of *Prrx1* was sufficient to speed up the response to Sonic Hedgehog agonist (SAG). Our results support the hypothesis that cell-to-cell phenotypic variation is in part caused by fluctuations in the activities of TFs.

## Results

### Variability in the HH response

We first asked whether clonal cells growing in the same environment show variability in their responses to HH signaling. To address this question, we used NIH3T3-CG cells which are an established model of HH signaling (Pusapati et al. 2018; Kinnebrew et al. 2019). Treating these cells with SAG activates the pathway, resulting in elevated activity of Gli transcription factors (Briscoe and Thérond 2013; Lee, Zhao, and Ingham 2016; J. H. Kong, Siebold, and Rohatgi 2019). This increased activity of Gli is read out by a genome-integrated GFP reporter gene regulated by eight Gli binding sites (Supplementary Figure 1). To assess variability in the HH response we treated NIH3T3-CG cells with SAG and monitored the response of the GFP reporter gene by flow cytometry (Figure 1A). Although monocultures of NIH3T3-CG cells are clonal and are grown in the same controlled environment, we detected significant cell-to-cell variability in reporter gene expression in response to SAG (Figure 1B). For example, at 30 hours post SAG treatment only 39% of cells had activated the reporter gene, whereas by 92 hours most cells responded (Supplementary Figure 2). We observed similar variability of response in multiple experimental replicates grown on different days and derived from multiple single-cell clones. What accounts for the difference between fast and slow responding cells in monocultures with no genetic or environmental variation?

**Figure 1:**
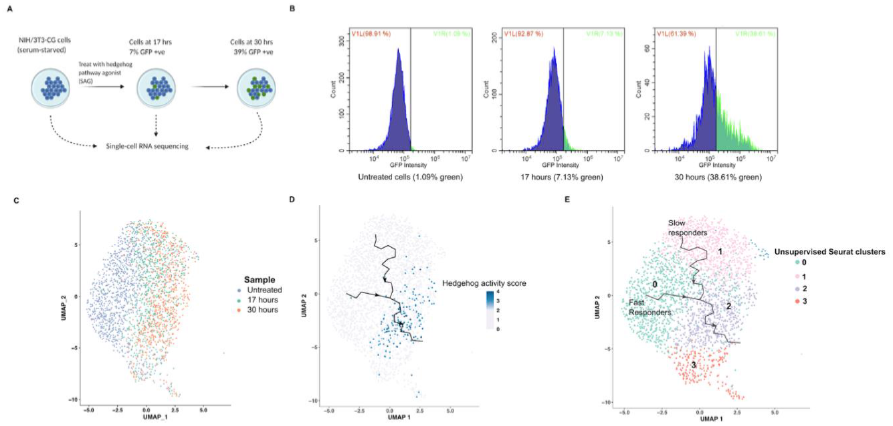
Cells show variability in response to hedgehog stimulation. (A) Cells were treated with hedgehog agonist SAG for 0, 17 and 30 hours and collected for single-cell RNA sequencing and flow cytometry. (B) Flow cytometry results at 17 and 30 hours indicate that cells show variability in their timing of response, not all same cells respond at the same time (C) Cells clustered with a clear temporal progression of hedgehog response when colored by sample time-point (D) Cells were scored for hedgehog pathway response by looking at mRNA of four canonical hedgehog response genes, responder cells cluster in one part of the graph. Response trajectories were inferred using trajectory analysis software (E) Unsupervised clustering of cells is shown. Cells take two different trajectories to respond to hedgehog pathway activity, one path is fast responding consisting of only cells at zero hours, the other response is from slow responders consisting of cells at 0, 17 and 30 hours.

### Fast and slow responding NIH3T3-CG cells

We asked whether cells that respond quickly to SAG derive from untreated cells with expression profiles that are distinct from the slow responding cells. To do this, we performed scRNA-seq on cells at 0, 17, and 30 hours after the addition of SAG. The SAG addition was done in a staggered manner to process all three scRNA-seq libraries in a single batch (Methods, Supplementary Figure 3). We first visualized the cells at the three time-points using UMAP (Figure 1C) and performed unsupervised clustering of cells. On the same UMAP plot, we colored the cells for Hedgehog response using four Hedgehog response genes (Methods) and observed a region of the UMAP where the responding cells reside (Figure 1D). We observed similar results when we visualized cells using Principal Components Analysis (PCA) (Supplementary Figure 4), but UMAP highlighted the local differences between cells better and captured all the variation in two dimensions.

Next, we used trajectory analysis (Qiu et al. 2017; Trapnell et al. 4/2014) to identify the cell states from which slow and fast responders derive. We observed two distinct trajectories after SAG treatment that lead towards cells expressing genes indicative of the Hedgehog response (Figure 1D). Fast responding cells start in cluster 0 and follow a trajectory that leads into cells that express Hedgehog responsive genes at 17 and 30 hours (Figure 1E). Slow responding cells start in cluster 1 and many of these cells remain in the same cluster at 17 and 30 hours, while other slow responding cells follow a trajectory that leads towards but does not reach Hedgehog responding cells. We interpret this result to mean that untreated NIH3T3-CG cells consist of two distinct subpopulations, one that is primed to respond quickly to SAG (fast-responders - cells in cluster 0 of Figure 1C) and another that takes longer to respond (slow responders - cells in cluster 1 of Figure 1C). Very few untreated cells lie outside these two clusters. The fact that these two subpopulations of untreated cells, identified through unsupervised clustering, are in distinct regions of the UMAP plot suggests that their global mRNA expression profiles define the fast and slow responding states.

We used a second trajectory analysis software, Slingshot (Street et al. 2018), to verify the trajectory inferred using Monocle. Both Monocle and Slingshot were both rated accurate trajectory inference tools in a review evaluating different trajectory analysis software (Saelens et al. 2019). We observed similar results with Slingshot (Supplementary Figure 5) as we did with Monocle; we find two different trajectories, one starting from the fast responders and another from the slow responders, leading to the hedgehog responsive cells.

One hypothesis that might explain cell-to-cell differences in the timing of the Hedgehog response between the two subpopulations would be transient activation of Hedgehog signaling in the absence of ligand. Cells that are in the midst of transiently activating the pathway would then respond faster to extracellular ligand than the remaining cells. To test this hypothesis, we looked for subsets of cells that expressed targets of Hedgehog signaling. In the unstimulated population we did not observe individual cells with activation of Hedgehog target genes (Supplementary Figure 6). Thus, Hedgehog signaling appears to be regulated tightly enough that stochastic activation of the pathway in the absence of ligand is unlikely to account for the cell-to-cell differences in cellular response after the addition of ligand.

### Differences in cell cycle state do not fully explain fast and slow responding cells

We next asked whether differences in the cell-cycle state of cells might explain the differences between the fast and slow responding cells. We used the global expression profiles of untreated cells to assign them to different phases of the cell cycle and asked whether fast responding cells were enriched for any specific phase of the cell-cycle. We observed a 1.5 fold enrichment (74.3% of fast responders vs 50.4% in the slow responders) of G1 cells in the fast responders compared to the slow responders (Supplementary Figure 7). While we found a statistically significant enrichment, the fact that half of slow responding cells were found in G1 suggests that being in G1 is not sufficient to respond faster. We conclude that cell-cycle differences do not fully explain the differences between fast and slow responding cells.

### Differentially expressed transcription factors and pathways in the fast-responder cells

We identified genes that are differentially expressed between untreated cells defined as either fast or slow responding (Methods). We focused on the top 300 genes, ranked by fold-change, that were statistically significantly different between the fast and slow populations (Supplementary Data 1). We use these 300 genes in subsequent analyses as markers of the fast responder state. Among the 300 genes, 31 genes are TFs; we decided to focus on the TFs in the differentially expressed genes because we hypothesize that fluctuations in TF activities account for the differences between fast and slow cells. The top differentially expressed TFs are shown in Supplementary Table 1.

We also asked if any biological pathways were among differentially expressed genes between the fast and slow responding cells through a Gene Ontology(GO) analysis (Methods). We observed that the cholesterol biosynthesis pathway is significantly enriched in the set of genes differentially expressed between the two groups (Supplementary Table 2, Enrichment = 28.38, p-value = 1.7e-4, n = 5 out of 13 genes). Cholesterol boosts Hedgehog signaling by serving as a cofactor for Smoothened (Huang et al. 2018, 2016; Kinnebrew et al. 2019; Luchetti et al. 2016; Radhakrishnan, Rohatgi, and Siebold 2020). Increased cholesterol biosynthesis may contribute to the phenotype of fast responding cells.

### Slow responding cells pass through the fast state before responding

Slow responding cells might respond more slowly because they first have to enter the fast responding state from their current state or because they follow a distinct set of cellular states that lead to the Hedgehog response. We distinguished between these two possibilities by examining the set of genes that come on in the slow responders at later time points. Specifically we examined cells from hour 17 and hour 30 in the slow-responding cluster (cluster 1 in Figure 1D) and asked what genes are differentially expressed compared to untreated cells in the same cluster. When the slower responders respond, we observe that they turn on the gene signature of the fast responding cells. At 17 hours these cells express 32 of the 300 fast responder genes (total number of DE genes = 117), and at 30 hours they express 48 of the 300 fast responder genes (total number of DE genes = 196), these are both statistically significant enrichments over random expectation (Hypergeometric test; p-values = 7.9e-27, 1.8e-37). We interpret this result to mean that the slower responders take longer to follow the same trajectory as fast responding cells rather than following a different trajectory.

### Overexpressing candidate transcription factors partially recreates the fast responder gene signature

Our results suggest that fluctuations in the activities of a small set of transcription factors and/or the upregulation of cholesterol biosynthesis pathway causes the differences between fast and slow responding cells. To test this hypothesis, we over-expressed these transcription factors and the cholesterol biosynthesis pathway and measured the bulk expression profiles of the resulting cells. We focused on the top three transcription factors based on fold-change in the fast responder cells (Supplementary Table 1): *Jun, Egr1 and Prrx1*. We also tested the transcription factor *Srebf2*, the main regulator of the cholesterol biosynthesis pathway (Horton et al. 1998), because this pathway is upregulated in the fast responder cells (Supplementary Table 2). We chose *Snai1* as a control transcription factor due to its lower fold change (fold-change 1.21). We designed plasmids encoding the coding regions of *Jun, Egr1, Prrx1, Srebf2 and Snai1* under the control of a strong CMV promoter (Supplementary Figures 8, 9). We used a plasmid containing an mCherry reporter gene driven by a CMV promoter as a transfection control plasmid. We transfected these plasmids into NIH3T3-CG cells and performed bulk RNA-seq. We identified genes that are differentially expressed before and after transfection of each of the transcription factors but not in the control transfection plasmid (Supplementary Data 2, 3, 4, 5, 6).

Overexpression of four of the five transcription factors (*Prrx1, Snai1, Jun and Srebf2*) individually misexpressed a significant number of the fast responder genes (Figure 2; Hypergeometric test: p-values = 9.7e-4, 6.3e-7, 1.2e-10, 1.9e-16), whereas *Egr1* overexpression did not (p-value = 0.13). We found that overexpressing *Snai1*, our control TF with a lower fold-change, also misexpressed a significant number of fast-responder genes indicating that even TFs with lower fold-changes might be involved in creating the fast-responder signature in cells. We found that the union of all four of these TFs (*Prrx1, Snai1, Jun and Srebf2*) resulted in 115 out of the 300 fast responder gene signature, this is smaller than the naive sum of the individual target genes and indicates overlap in the sets of target genes regulated by the TFs. Overall, we interpret this result to mean that each TF independently regulates a subset of the fast-responder signature.

**Figure 2:**
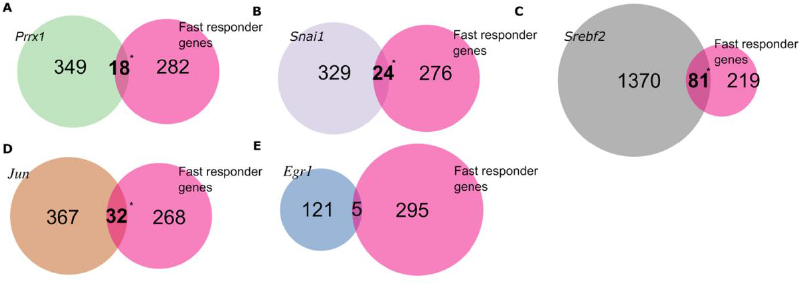
Overexpressing transcription factors partially recreates the fast responder gene expression signature. We overexpressed six transcription factors, one at a time, *Prrx1* (A), *Snai1* (B), *Srebf2* (C), *Jun* (D) and *Egr1* (E) using a plasmid transfection assay followed by bulk RNA sequencing. We computed the overlap between the genes that change expression upon transcription factor overexpression and the 300 genes in the fast-responder gene signature identified using the single-cell data.

### *Prrx1* is a regulator of the fast responder state

We next asked whether the overexpression of any of these transcription factors was sufficient to increase the fraction of cells that respond to the Hedgehog signaling pathway. To test this prediction we engineered cells, using lentiviral vectors (Methods, Supplementary Figures 10, 11), carrying Dox-inducible(Supplementary Figure 12) versions of *Prrx1, Srebf2* and *Snai1* integrated into their genomes. This allowed us to compare the fraction of cells that respond to SAG when a TF is either induced or uninduced.

Inducing *Prrx1* expression resulted in more cells responding to the Hedgehog pathway (Figure 3A, Supplementary Figure 13). In the +Dox condition, at 32 hours post SAG treatment, 87.9% of cells become GFP+, whereas in the -Dox condition 65.4% of cells become GFP+. We did not observe a difference between the two groups with the induction of *Snai1*. Strong induction of *Srebf2* was toxic to cells, and at induction levels that were not toxic we did not observe an increase in the response to SAG (Supplementary Figures 14 and 15). From these results, we conclude that *Prrx1* plays a role in the regulation of the Hedgehog response.

**Figure 3:**
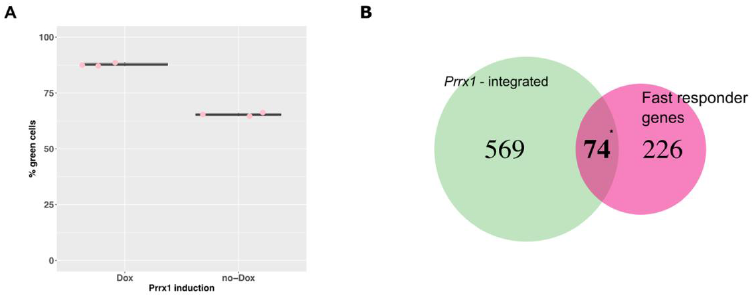
Inducing *Prrx1* expression makes more cells fast-responders. (A) We induced *Prrx1* expression in engineered cells by adding Doxycycline prior to performing the hedgehog assay. We compared the hedgehog response of induced cells to cells that were not induced using flow cytometry 32 hours post SAG treatment. (B) We identified the genes that change expression when *Prrx1* is overexpressed by bulk RNA sequencing. We then computed the overlap of these genes with the 300 genes in the fast responder signature.

Bulk RNA-seq profiles from *Prrx1* induced cells (Supplementary Data 7) revealed a stronger overlap with fast responding genes than in the plasmid overexpressed cells (Figure 3B; n = 74 out of 300, p-value = 4e-34, enrichment = 5.37 fold). This stronger measured response is likely because in transiently transfected cells only a subset of the cells are expressing *Prrx1* whereas in the lentiviral transduced cells all the cells in the population are misexpressing *Prrx1*. Induction of *Prrx1* results in significant enrichment of genes involved in the cholesterol biosynthesis pathway (adjusted p-value = 0.025, enrichment = 10.42, n = 4 out of 13 genes in the pathway). *Prrx1* induction also results in the upregulation of *Gli2*, the primary effector TF of Hedgehog signaling (J. H. Kong, Siebold, and Rohatgi 2019; Briscoe and Thérond 2013; Lee, Zhao, and Ingham 2016). *Gli2* is regulated post-translationally by Hedgehog signaling, so the upregulation of *Gli2* in uninduced cells may result in a larger pool of *Gli2* protein to activate when the pathway is activated. Our results indicate that misexpression of *Prrx1* can recreate a substantial part of responder signature, which we interpret as the reason more of these cells respond to Hedgehog signaling.

### Most cells activate the fast responder gene signature upon *Prrx1* induction

We next examined the trajectory taken by cells overexpressing *Prrx1*. We generated scRNA-seq profiles of *Prrx1*-induced and uninduced cells followed by stimulation with SAG (Figure 4A). We started with the cells carrying the genome-integrated inducible *Prrx1* cassette described in the previous section. We grew cells in the absence or presence of Dox, to induce *Prrx1* expression, and measured the HH response by flow cytometry. We again observed a larger response in *Prrx1*-induced cells than in uninduced cells (Supplementary Figure 16). We then collected cells at three time points after SAG treatment for both *Prrx1*-induced and uninduced cells and performed scRNA-seq.

**Figure 4:**
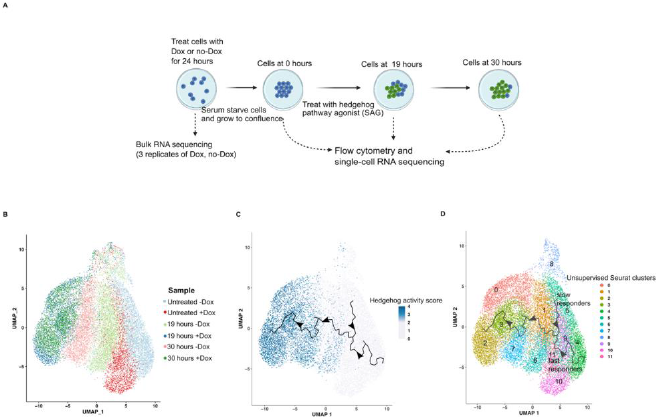
*Prrx1* induction results in faster response trajectory. (A) Cells were grown in the absence and presence of Doxycycline for 24 hours, to induce *Prrx1* expression, and treated with the hedgehog pathway agonist for 0 (Untreated), 19 and 30 hours. (B) The clustering of the cells from the three time points across the two conditions (C) The inferred trajectory of hedgehog response. Cells were scored and colored for hedgehog pathway response by looking at mRNA expression of four canonical hedgehog response genes (D) The unsupervised clustering of the cells using Seurat. Fast responders and slow responders predicted using a cell-type classifier are shown.

*Prrx1* induced cells follow a faster and stronger response trajectory compared to their uninduced counterparts. When *Prrx1* is not induced, cells from each time-point again cluster separately, suggesting a steady progression of response to SAG through time (Figure 4B). By contrast, when we induced *Prrx1* expression with Dox, cells at the 19 hr and 30 hr time point cluster together, suggesting a faster response to SAG induction (Figure 4B). Overlaying the Hedgehog response on these clusters supports this interpretation as *Prrx1*-induced cells at 19 hours show similar levels of the Hedgehog response as uninduced cells at 30 hrs (Figure 4C). However, even at 30 hrs induced and uninduced cells do not cluster together, which suggests that Prrx1 induction results in a stronger response to Hedgehog in cells that do respond. Taken together, these results show that the induction of *Prrx1* causes more cells to respond to SAG, and that those cells that do respond, respond faster and stronger than cells that do not express *Prrx1*.

The trajectory analysis also revealed differences between induced and uninduced cells in the early response to SAG that are consistent with the idea that *Prrx1* causes cells to adopt fast-responding expression profiles. Uninduced cells again showed distinct converging trajectories from slow and fast responding states (clusters 4, 5 in Figure 4D), but Prrx1-induced cells respond primarily along the fast trajectory. In addition, the Prrx1-induced cells start from a position along the fast trajectory that is closer to the later time points than the uninduced cells (clusters 9, 10, 11 in Figure 4D), which suggests that they are “further along” the fast trajectory than uninduced cells even before stimulation with SAG.

We next asked whether inducing Prrx1 creates more fast-responding cells. To detect fast responding cells in this experiment we used a model-based cell-type classifier Garnett (Pliner, Shendure, and Trapnell 2019). We trained the classifier to learn the features of fast-responding and slow responding cells using the cells from the untreated condition shown in Figure 1. We then used the classifier to classify untreated cells grown in the +Dox and -Dox conditions as fast responders and slow responders. In the untreated cells grown in -Dox condition, the classifier classified 19% of cells as fast-responders, 48% as slow responders and 33% of the cells as unknown. The fast responders in this group separate from the slow responders in the UMAP plots and are slightly ahead in the response trajectory even before SAG treatment (Supplementary Figure 17 A, B). In contrast, in the untreated cells from the +Dox condition the classifier classified 91% of cells as fast-responders, 2% slow-responders and 7% as unknown (Supplementary Figure 17 C, D). Thus in the presence of Dox, the majority of cells display the fast-responder expression signature. We infer from this result that inducing *Prrx1* generates a fast responder cell-state which makes more cells respond to the Hedgehog agonist, as early as 19 hours.

## Discussion

We hypothesized that fluctuations in the activities of TFs produce transient changes in gene expression that can generate phenotypic differences among clonal cells in the same environment. Consistent with this hypothesis, we showed that distinct gene expression profiles define fast and slow responding states among clonal NIH3T3-CG cells, and that the activities of a small number of TFs accounts for a substantial fraction of the expression differences that define the two states. The idea that these expression profiles cause differences in the response to Hedgehog signaling is supported by the observation that misexpression of *Prrx1* is sufficient to generate part of the fast responding expression signature and drive a faster and stronger response to Hedgehog in a larger fraction of cells. However, because *Prrx1* only accounts for part of the fast responding expression signature, there must be fluctuations in other TFs(Sigal et al. 11/2006), or other types of signaling molecules, that generate the fast responding cell state.

Much deeper sequencing of unstimulated cells might reveal groups of differentially expressed target genes that could indicate which TFs and signaling molecules are involved, much the same way in which coherent differential expression of the cholesterol biosynthesis pathway suggested the involvement of *Srebf2* in the fast responding state. Alternatively, if the transient activation of non-TFs (e.g. receptors, kinases)(Colman-Lerner et al. 2005) underlies the fast responding state, then it will be necessary to identify specific changes in gene expression that are diagnostic of changes in the activities of such genes.

We observe parallels between our model underlying variability in the Hedgehog response and other previously described systems where fluctuations of a few key molecules underlie cell-to-cell variability in phenotypic outcomes. For example, differences in the levels of a few apoptotic regulator proteins prior to drug treatment determines how fast cells die in the presence of an apoptosis inducing ligand (Spencer et al. 2009). Fluctuations of a few resistance genes in cancer cells results in cell states that are resistant to cancer drugs, (Emert et al. 2021; Shaffer et al. 2020) though the exact mechanisms underlying the emergence of such cell states remains unknown and is hypothesized to involve the fluctuation of multiple upstream transcription factors. In stem cells, fluctuations of a key transcription factor Nanog affect whether cells differentiate or remain in the pluripotent state (Miyanari and Torres-Padilla 2012; Torres-Padilla and Chambers 2014). A recent study using fluorescence microscopy determined that most of the variability in the JAK-STAT signaling response can be attributed to fluctuations in the molecular content of cells (Topolewski et al. 2022). Single-cell transcriptomic approaches, such as those used in this study, can shed light on the specific molecular differences between cells that result in heterogeneity of response.

Our study also has important implications for improving the efficiency of cellular assays. For example, reprogramming assays can be made more efficient if we understand why some cells are successfully reprogrammed while other cells result in ‘dead-end’ states (Biddy et al. 2018; Graf and Enver 2009; Francesconi et al. 2019). Fluctuations of transcription factors in cells prior to reprogramming might underlie the heterogenous outcome. For example, in hematopoietic progenitor cells, levels of a stem cell marker determine which lineage a cell differentiates towards (Chang et al. 2008). Isolating cells with different levels of such proteins may improve reprogramming efficiency.

We have inferred the Hedgehog response trajectories using computational methods that look at gene expression similarities between single-cells. A complementary measurement of the trajectory would involve using transcribing molecular barcodes to tag the cells prior to single-cell RNA sequencing (W. Kong et al. 2020). This approach could reveal how fast cells transition between the slow and fast responder states (Hormoz et al. 11/2016; Stumpf et al. 2017; Larsson et al. 2021). We have used overexpression experiments to infer the role of *Prrx1* in the fast response. Using GFP fused versions of *Prrx1* would help separate the fast responder population from the slow responders and perform functional assays on these subsets of cells. Finally, we have used a widely used cell-culture model of Hedgehog signaling, and how our findings apply to *in vivo* Hedgehog signaling remains to be tested.

One future direction of this work will be to determine whether the fast responding state is specific for Hedgehog signaling or if this state makes cells differentially responsive to other signaling pathways and perturbations. We speculate that similar variability in the activities of different TFs may underlie other phenotypic differences among genetically identical cells and experimental approaches similar to ours can be used to uncover this variability.

## Materials and Methods

### Cell Culture

We grew NIH/3T3-CG cells and their derivatives in DMEM media supplemented with sodium pyruvate, 10% Bovine Serum (Gibco 16170078) and 1% Penicillin-Streptomycin. We grew the cells in an incubator maintained at 37 degrees celsius with 5% CO2.

### RNA Sequencing Experiments

We generated single-cell RNA sequencing libraries using the 10X Single Cell 3’ Reagent Kits v3.1 (10X genomics 1000269). We first released the cells from the cell culture flasks by adding Trypsin 0.25%. We then prepared a cell suspension by following manufacturer’s instructions and targeting a final capture of 2000 cells for every sample. We then proceeded with library construction as described in the 10X protocol. We sequenced the final libraries at the DSIL and MGI at Washington University in St. Louis. Bulk RNA sequencing was performed using the services of Novogene corporation. Briefly, we extracted total RNA from cells, performed initial QC and shipped it to Novogene where library preparation and sequencing were performed targeting 20 million reads per sample.

### RNA sequencing analysis

We obtained counts for each transcript in each cell by using Cell Ranger version 3.1.0 (https://support.10xgenomics.com/single-cell-gene-expression/software/pipelines/latest/using/count). We analyzed the cell by count matrix using the standard analysis workflow of Seurat version 3.1.5.(Butler et al. 2018; Stuart et al. 2019) For quality control, we kept cells that had at-least 10,000 UMIs per cell, had < 20 % of UMIs mapped to mitochondrial genes, and had between 2000 and 8000 genes covered. After QC, for the initial single-cell sequencing experiment we were left with transcriptome measurements for 1,192 cells for the untreated time-point, 797 cells at 17 hours and 899 cells at 30 hours. For the second single-cell RNA-sequencing experiment using the inducible Prrx1 cell-line, under the no-Dox condition we were left with 4532 cells at the untreated time-point, 3726 cells at 19 hours and 2645 cells at 30 hours after QC. Under the Dox condition, after QC we were left with 2362 cells at the untreated time-point, 3014 cells at 19 hours and 2645 cells at 30 hours.

We performed dimension reduction of single cells using UMAP(McInnes, Healy, and Melville 2018). We identified differentially expressed genes between clusters using the FindMarkers function in Seurat. We performed trajectory analysis using Monocle3 version 1.0.0(Trapnell et al. 4/2014; Qiu et al. 2017) and Slingshot version 1.4.0 (Street et al. 2018). Details regarding the parameters used for the various programs are available in the R notebooks made available in the Zenodo repository listed below. We identified pathways that were enriched in the set of differentially expressed genes using the Gene Ontology web resource(Ashburner et al. 2000; Mi et al. 2019; Gene Ontology Consortium 2021).

We computed the hedgehog pathway activity score for each cell by looking at the mRNA expression of four genes - the GFP reporter, *Gli1, Gli2* and *Ptch1*. For each of the four genes in each cell, we assigned a score of 1 if the gene is detected above background expression level and 0 if not. We then sum over all four genes to obtain a score in the range 0-4 for each cell.

For bulk RNA-sequencing analysis we pseudo-aligned the reads to the mouse reference cDNA using Kallisto version 0.43.0 (Bray et al. 2016). We used the counts from Kallisto to identify differentially expressed genes using DESEQ2 version 1.26.0 (Love, Huber, and Anders 2014). We used a false-discovery rate of 0.05 to annotate genes as differentially expressed across conditions.

### Hedgehog Assay

We performed the Hedgehog assay on NIH/3T3-CG cells as previously reported in Pusapati et. al. Briefly, we grew cells to confluence in media containing regular amounts (10%) of serum. We then switched the cells to low serum media (0.5% serum) overnight, and treated the cells with the Hedgehog pathway agonist SAG (Tocris Biosciences #4366) at 100 nM concentration. We measured the amount of fluorescence in the cells on a flow cytometer after allowing for the appropriate time period of response depending on the experiment. For the initial experiment in Figure 1 we collected cells at 17 hours and 30 hours. For the Prrx1 overexpression single-cell experiment we collected cells at 19 hours and 30 hours. For both these experiments, we treated cells that didn’t receive SAG as untreated cells or time-point zero. The time course treatments were done in a staggered manner so that all the cells could be harvested at the same time for single-cell RNA sequencing library preparation. This enabled us to process all the cells in one batch to minimize experimental variability for single-cell RNA sequencing.

### Flow Cytometry

We performed the flow cytometry experiments on a Beckman Coulter Cytoflex S instrument. We performed QC based on manufacturer provided QC beads prior to every experiment. We used the same cytometer gain settings for all experiments. To prepare cells for flow cytometry, we first released cells from culture wells by adding Trypsin and then added appropriate volume of culture media to neutralize Trypsin. We gated cells based on forward scatter and side scatter and measured GFP intensity on the FITC channel. We used a negative control sample (no SAG added) to identify the cutoff for the GFP intensity and measured the proportion of responders as the percentage of cells above this cutoff under different experimental conditions.

### Plasmid Transfection

To transfect the plasmids overexpressing the transcription factors we used Lipofectamine 3000 (Invitrogen L3000001) reagent. We used 2.5 ug of DNA per transfection reaction, on a six-well plate, and followed manufacturers instructions for amounts of Lipofectamine and P3000 reagents. We measured transfection efficiency on a flow cytometer using the fluorescence of a reporter gene on the transfected plasmid or a control plasmid transfected in parallel. If we observed sufficient transfection efficiency (> 50% of cells express transfected plasmid) we extracted total RNA, 24 hours post transfection, from the cells and performed bulk RNA sequencing. The control plasmid used is a mCherry reporter gene driven by a CMV promoter.

### Lentiviral generation and transduction

We chose the canonical coding transcripts and sequences for all the genes from UniProt(“UniProt: The Universal Protein Knowledgebase in 2021” 2021). We then cloned the protein coding regions of *Prrx1, Snai1* and *Srebf2* to the pINDUCER21 Dox-inducible lentiviral vector (Meerbrey et al. 2011) using the services of Genscript (Addgene #46948, Supplementary Figures 6 and 7). We used the pINDUCER21 system since it allows us to control the level of over-expression of a gene with precision by adding different levels of Doxycycline to the cell growth media. This construct also has a fluorophore (miRFP670) on it which allows us to isolate cells that contain the integrated construct. The Hope Center at Washington University in St. Louis generated high-titer lentiviruses using the constructs. Detailed lentivirus generation protocol used by the Center is described in their publication (Li et al. 2012).

We used the generated virus to transduce the NIH3T3-CG cells and used 4 ug/ml polybrene to maximize transduction efficiency. After 24 hours of cell-growth in the transduction media, we replaced the media with fresh regular media. We used a fluorescent marker of integration to sort single-cells (miRFP670) using a Sony SH800 cell-sorter and grew out single-cell clones that we evaluated for TF induction.

We evaluated single-cell clones based on induction levels assessed by qPCR. For induction of clones we used Doxycycline (Dox) at a concentration of 500 ng/ml. For transcription factor induction followed by RNA-seq experiments, cells were treated with Dox for a period of 24 hours to induce TF expression. We identified one clone for each transcription factor that offered a good level of inducibility (Supplementary Figure 8). We then used these clonal cell lines as a model to study the effect of TF overexpression on the Hedgehog assay.

To perform the Hedgehog assay, we grew the cells in two different growth conditions - in one condition we added Dox to the media and in the other condition we omitted Dox. We then grew the cells to confluence, for 24 hours, and added SAG to both populations of cells to initiate the Hedgehog pathway response. We used flow cytometry to determine the effect of the overexpression of these transcription factors on the response to Hedgehog stimulation by measuring the GFP fluorescence of the reporter gene on the FITC channel.

### qPCR Run and Analysis

We first extracted total RNA from cells using the Qiagen RNEasy kit(Qiagen 74004). We generated cDNA from the total RNA using the RDRT reagent (Sigma RDRT-100RXN) by following manufacturer’s protocol. We then mixed SYBR green PCR master mix(Applied Biosystems 4301955) with 2 ul of cDNA from the previous step, water and primers to set up a standard qPCR run on the QuantStudio instrument (Applied Biosystems). For the transcription factor induction experiments using the transduced cell-lines, we used the no-Dox sample as the baseline for computing the delta Ct value. We used HPRT primers to normalize as a within sample control for TF expression (Supplementary Table 3). We analyzed the results of the QuantStudio run using the Design and Analysis 2 software from Thermo Fisher.

## Acknowledgements

NIH3T3-CG cells were a gift from the Rajat Rohatgi lab at Stanford University. We thank Ganesh Pusapati, Maia Kinnebrew and Siggy Nachtergaele for protocols and advice setting up the Hedgehog assay on NIH3T3-CG cells. We thank the members of the Cohen lab for a critical reading of the manuscript. We thank Xuhua Chen for help with the 10X protocol. We thank Abul Usmani for advice regarding RNA sequencing protocols. We thank Jessica Hoisington-Lopez and ML Crosby in the DNA Sequencing Innovation Lab (DSIL) for assistance with high-throughput sequencing. We thank Mingjie Li and the Hope Center at Washington University in St. Louis for generating lentiviruses.

## Author contributions

A.R and B.A.C. conceptualized and designed the project. A.R designed and performed all experiments and analyses. A.R and B.A.C. wrote the manuscript.

## Data Availability

The raw single-cell and bulk RNA sequencing data from this publication are available from GEO under the accession numbers GSE203134 and GSE206154. Analysis notebooks used for the analysis of single-cell data are available for download at Zenodo: https://doi.org/10.5281/zenodo.6981764

## Supplementary Information

### List of Supplementary Figures

Supplementary Figure 1 - Schematic of hedgehog pathway activation and GFP reporter

Supplementary Figure 2 - Most cells are responders by 72 hours

Supplementary Figure 3 - No evidence for batch effect

Supplementary Figure 4 - PCA visualizations are similar to UMAP

Supplementary Figure 5 - Slingshot trajectory analysis results

Supplementary Figure 6 - Hedgehog score distribution at the 0, 17 and 30 hour time-points

Supplementary Figure 7 - Cell cycle assignments for fast responders and slow responders

Supplementary Figure 8 - Plasmid design for Srebf2 transfection

Supplementary Figure 9 - Plasmid design for Snai1, Prrx1, Jun and Egr1 transfection

Supplementary Figure 10 - Plasmid design for Snai1, Prrx1 lentiviral transduction

Supplementary Figure 11 - Plasmid design for Srebf2 lentiviral transduction

Supplementary Figure 12 - qPCR induction plots for Prrx1, Srebf2 and Snai1

Supplementary Figure 13 - Raw data for the Prrx1 induced and uninduced cells shown in Figure 3A

Supplementary Figure 14 - no difference in percent green for Snai1 induction followed by SAG treatment

Supplementary Figure 15 - no difference in percent green for Srebf2 induction followed by SAG treatment

Supplementary Figure 16 - Cytometry results for cells used in Prrx1 induction single-cell RNA sequencing experiment

Supplementary Figure 17 - Predicted slow and fast responder cells using Garnett in the Prrx1 +/- Dox untreated cells

## List of Supplementary Data

Table S1 - Fast-responder gene signature - top 300 genes differentially expressed in the fast-responder cells compared to the slow-responder cells

Table S2 - Bulk RNA-seq Differential gene expression results - *Prrx1* transfection compared to mCherry control transfection

Table S3 - Bulk RNA-seq Differential gene expression results - *Snai1* transfection compared to mCherry control transfection

Table S4 - Bulk RNA-seq Differential gene expression results - *Srebf2* transfection compared to mCherry control transfection

Table S5 - Bulk RNA-seq Differential gene expression results - *Jun* transfection compared to mCherry control transfection

Table S6 - Bulk RNA-seq Differential gene expression results - *Egr1* transfection compared to mCherry control transfection

Table S7 - Bulk RNA-seq Differential gene expression results - *Prrx1* induction compared to no-Dox control

## Supplementary Figures

**Supplementary Figure 1:**
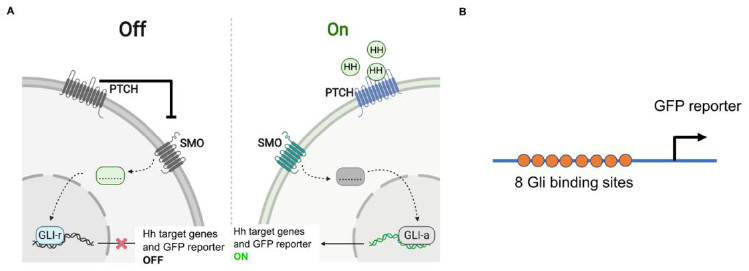
A simplified view of the Hedgehog pathway and activation of the GFP reporter. (A) In the absence of Hedgehog or SAG the pathway is turned off within a cell, the repression of SMO by PTCH leads to the transcription factor GLI being in its repressed form and it does not turn on the GFP reporter gene. In the presence of Hedgehog or SAG, PTCH no longer represses SMO, GLI acts as an activator and turns on the GFP reporter gene. (B) The cells used in our experiment were generated by *Pusapati et al*. and contain an integrated GFP reporter gene downstream of eight binding sites of GLI.

**Supplementary Figure 2:**
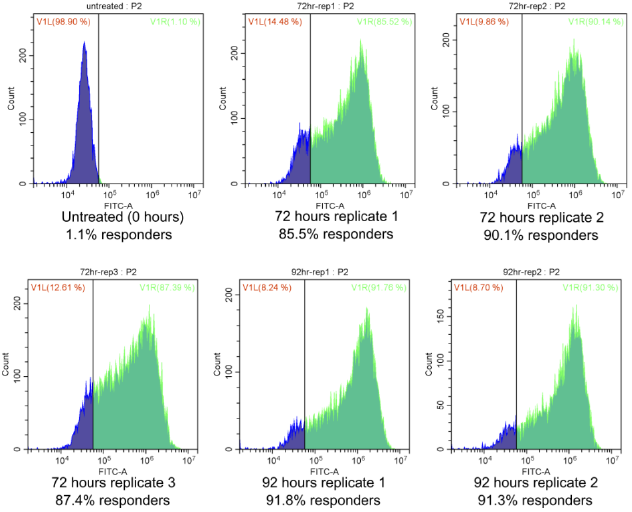
Flow cytometry results, at 0 (Untreated), 72 (3 biological replicates) and 92 hours (2 biological replicates) post SAG treatment. At 72 hours there are a mean of 87.6% responders across replicates and at 92 hours the mean fraction of responding cells is 91.6%

**Supplementary Figure 3:**
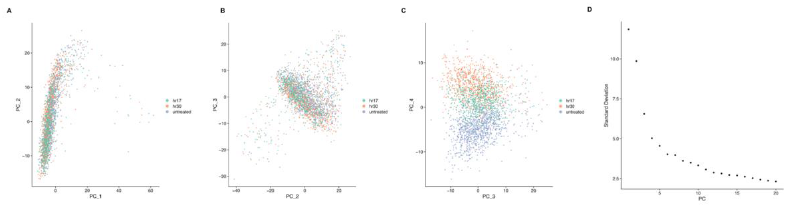
We see no evidence for batch effect since all the three samples were processed in a single batch. We plotted principal components for all the cells shown in Figure 1. Only PC4 (C) separates out the samples, and this separation is expected based on genes that turn on and off in a temporal manner in response to hedgehog. (D) shows the standard deviation explained by each principal component.

**Supplementary Figure 4:**
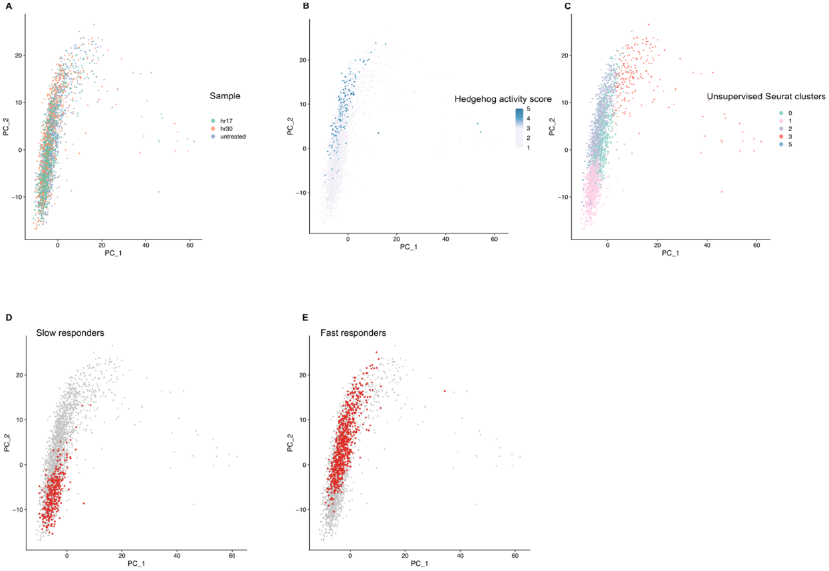
Visualizing cells with PCA shows similar results as visualizing with UMAP. We performed PCA on the cell by gene matrix and plotted the first two principal components for each cell. We colored cells by sample (A), hedgehog activity score (B) and unsupervised Seurat clusters (C). We highlighted the slow responders (D) and fast responders (E) identified in Figure 1 on the PCA plot and see that they separate out on the principal components plot.

**Supplementary Figure 5:**
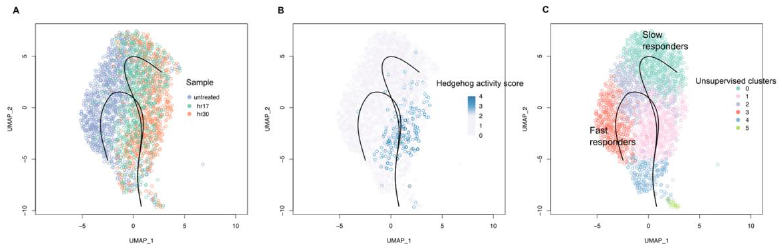
We used a second trajectory analysis tool Slingshot. Similar to results from Monocle in Figure 1, cells take two different trajectories in their response to SAG. We colored the cells by sample (A), hedgehog activity score (B) and unsupervised Seurat clusters (C) as in Figure 1.

**Supplementary Figure 6:**
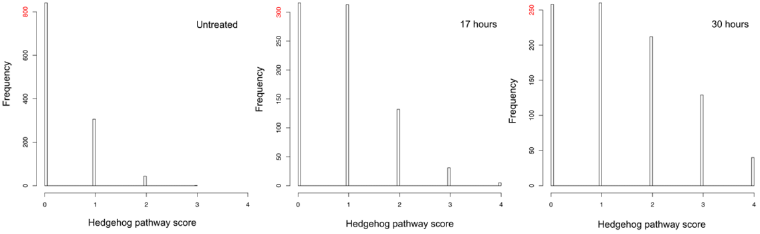
Hedgehog score distribution of cells at three different timepoints. For each cell, using the single-cell RNA sequencing data, we calculated the hedgehog score by looking at the expression of *Gli1, Gli2, Ptch1 and GFP*. We assigned each gene in each cell a score of 1 if detected above background expression level, and we summed across the four genes to compute a score in the range [0-4]. In the untreated population scores are close to zero, by 30 hours more cells respond and the responders are the cells with score >= 3. Note the y-axis for each panel is different, however the interpretation is made by looking at the distribution of scores within each panel.

**Supplementary Figure 7:**
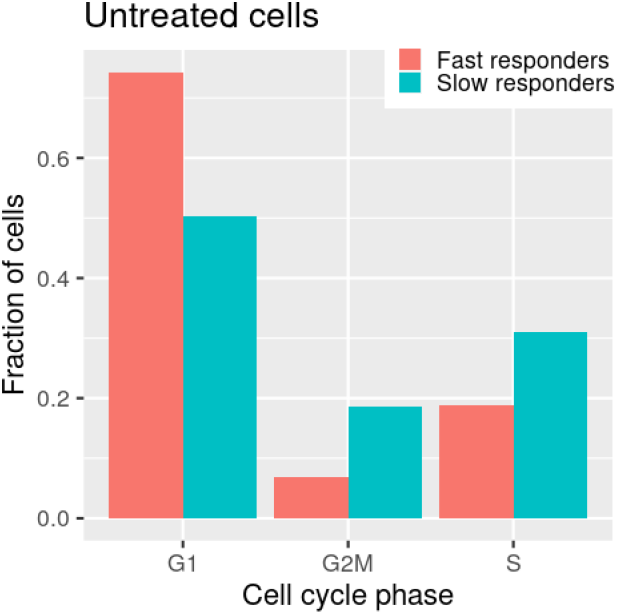
Inferred cell cycle phase for fast responders and slow responders. We inferred the cell cycle phase using the CellCycleScoring function in Seurat.

**Supplementary Figure 8:**
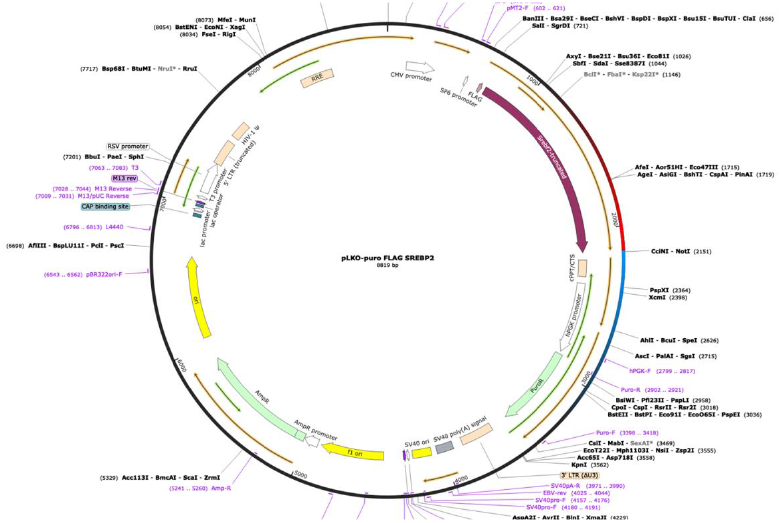
Plasmid design for overexpressing Srebf2 via transfection. The coding region of truncated *Srebf2* is under the control of the CMV promoter.

**Supplementary Figure 9:**
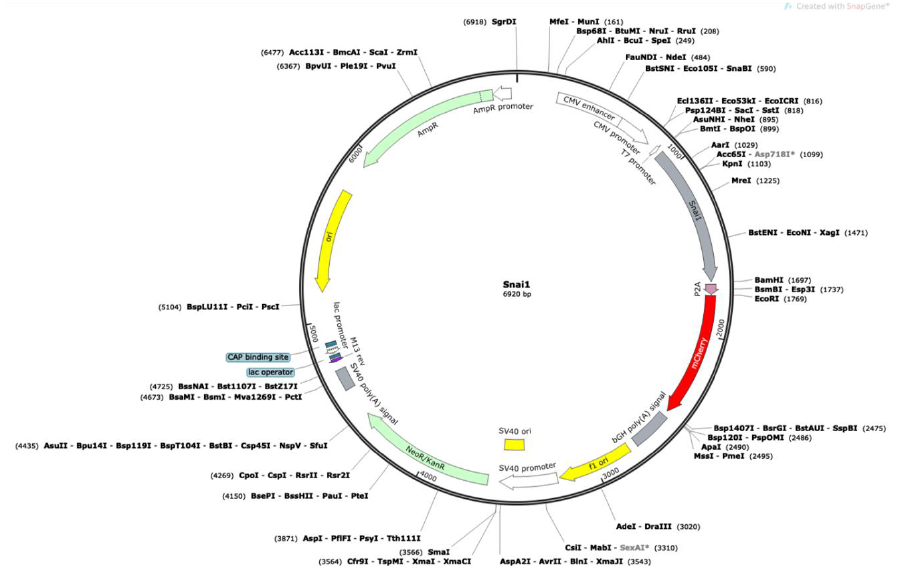
Plasmid design for overexpressing *Snai1*. The coding region of *Snai1* is regulated by the CMV promoter. This plasmid also contains a mCherry coding region to measure transfection efficiency on a cytometer. We used similarly designed plasmids for transfecting *Jun, Egr1* and *Prrx1* by replacing with the respective coding sequence.

**Supplementary Figure 10:**
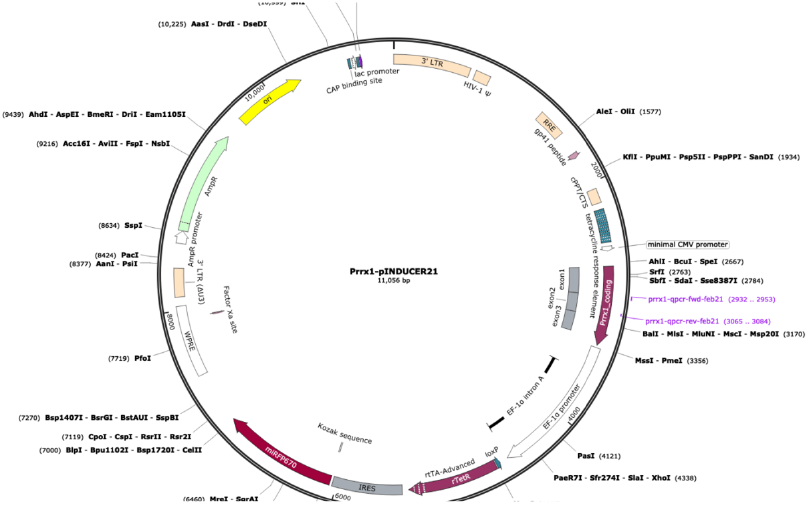
Construct for *Prrx1* pInducer induction. Lentiviruses were generated using this plasmid. We used the same design for *Snai1* by replacing the coding region of *Prrx1* with the coding region of *Snai1*.

**Supplementary Figure 11:**
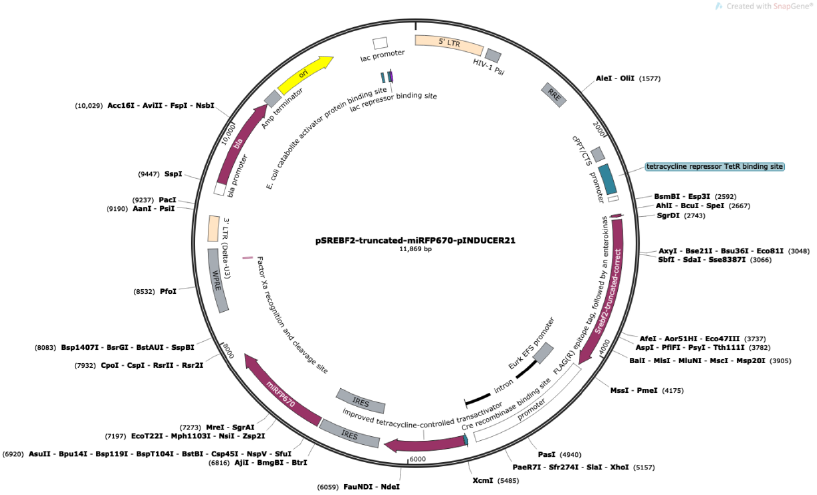
Construct for *Srebf2* pInducer induction. Lentiviruses were generated using this plasmid.

**Supplementary Figure 12:**
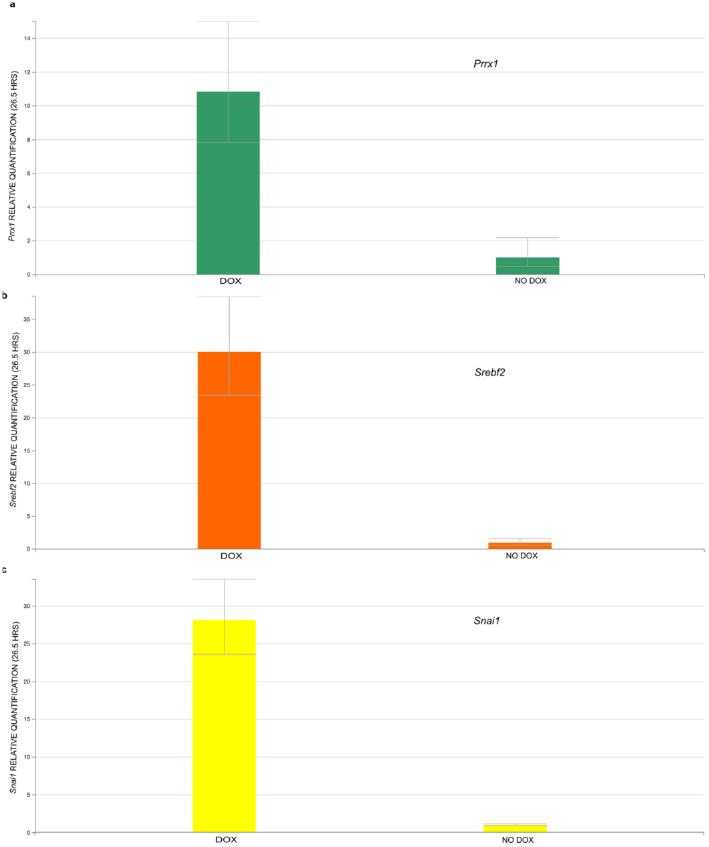
Induction of *Prrx1, Srebf2* and *Snai1*. Cell lines with inducible *Prrx1*(a), *Srebf2* (b) and *Snai1* (c) were treated with Doxycycline to induce the expression of the three genes. After 26.5 hours, we harvested the cells and used the RNA for qPCR assays. We used cells with no Doxycycline added as controls. We compared RNA levels in each condition to the housekeeping control gene *Hprt*.

**Supplementary Figure 13:**
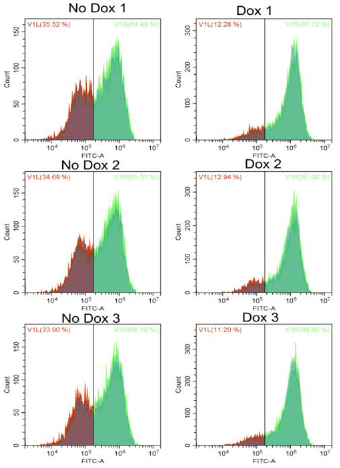
Raw cytometer plots for the data summarized in Figure 3A. Prrx1 inducible cells were grown in the presence and absence of Dox, treated with SAG and run on a cytometer at 32 hours. Three biological replicates were used for each condition.

**Supplementary Figure 14:**
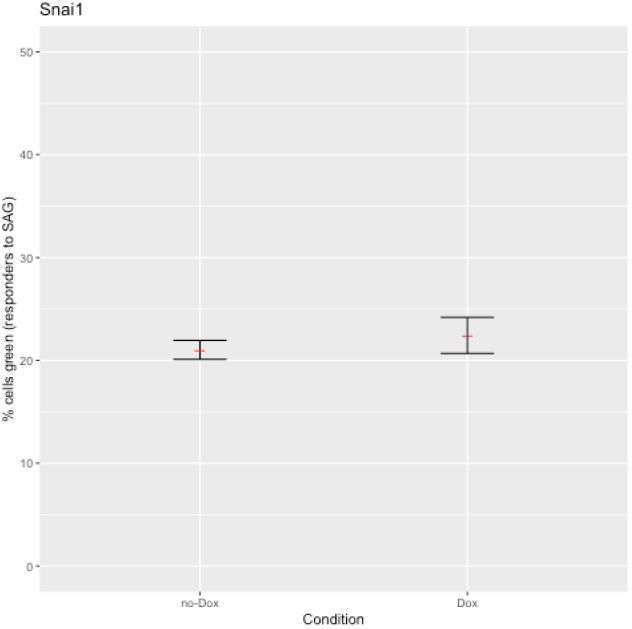
No difference in hedgehog assay response after *Snai1* induction. We grew cells integrated with inducible *Snai1* in the presence and absence of Doxycycline. We performed the Hedgehog assay on the two groups (three replicates each) of cells by adding SAG and observing the percent of cells that respond on the flow cytometer after 32 hours. Shown are the mean percentage of responders (in red) +/- standard error of the mean.

**Supplementary Figure 15:**
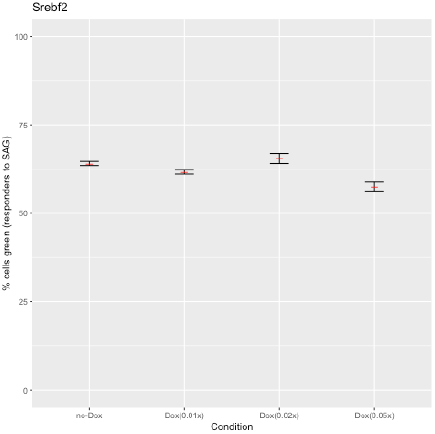
No difference in hedgehog assay response after *Srebf2* induction. We grew cells integrated with inducible *Srebf2* in the presence and absence of Doxycycline. We used three different concentrations of Dox. Cells did not survive in the presence of 1X and 0.1X Dox. In this figure, 1X Dox is the same concentration of Dox used for Prrx1 and Snai1 (500 ng/ml). We performed the hedgehog assay on the two groups (three technical replicates each) of cells by adding SAG and observing the percent of cells that respond on the flow cytometer after 46 hours. Shown are the mean percentage responders (in red) +/- standard error of the mean.

**Supplementary Figure 16:**
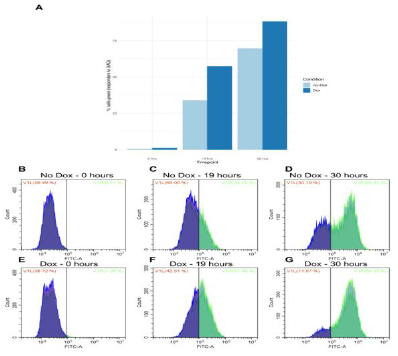
Response of Prrx1 induced and induced cells from the same pool as the cells used for the single-cell RNA sequencing experiments in Figure 4 (A) Percentage responding cells(GFP fluorescence) on a cytometer summarized at 0 (untreated), 19 and 30 hours. (B), (C), (D)Raw cytometer plots for the same cells in (A) in the no Dox condition. (E), (F), (G) Raw cytometer plots for the same cells in (A) grown in the Dox condition. This figure show one replicate at each condition used in the single-cell RNA sequencing experiment, biological replicates of *Prrx1*-induction response result is shown in Figure 3 and Supplementary Figure 13.

**Supplementary Figure 17:**
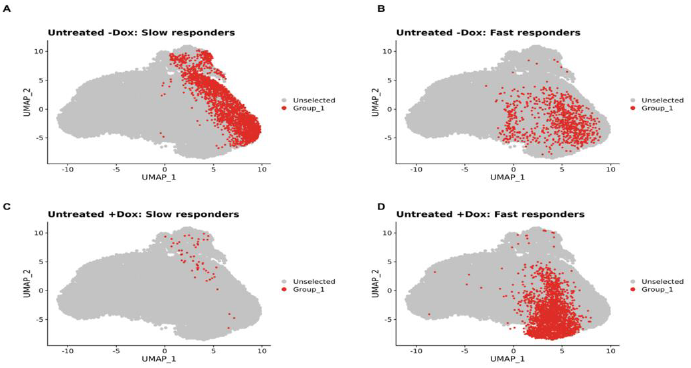
Predicted fast and slow responders from Garnett. We trained a cell-type classifier on the fast and slow responder cells in Figure 1 to learn the features of fast and slow responders. We next applied this trained classifier on untreated (0 hours) cells with inducible Prrx1, used in Figure 4, to identify the fast and slow responders in this population. (A) and (B) show the slow and fast responders identified by the model in the untreated cells grown in the absence of Dox. In the cells grown in the absence of Dox, the classifier identifies a mixture of slow and fast responders. © and (D) show the slow and fast responders identified by the model in the untreated cells grown in the presence of Dox. In the cells grown in the presence of Dox, most of the cells are classified as fast reponders by the classifier.

## Supplementary Tables

**Supplementary Table 1:** List of top transcription factors overexpressed in the fast responders.

| Transcription Factor | Fold-change (fast responders vs slow responders) |
| --- | --- |
| *Jun* | 1.55 |
| *Egr1* | 1.41 |
| *Prrx1* | 1.39 |

**Supplementary Table 2:** List of signaling pathways enriched in the fast responder cells compared to slow responders.

| Pathway | Genes in fast responder gene signature and in the pathway (Total number of genes in pathway) | Enrichment over random expectation (FDR adjusted p-value) |
| --- | --- | --- |
| Cholesterol biosynthesis | 5 (13) | 28.38 (1.7e-4) |
| Integrin signaling pathway | 20 (189) | 7.81 (6.2e-10) |
| Ubiquitin proteasome pathway | 5 (65) | 5.68 (4.1e-2) |
| Cytoskeletal regulation by Rho GTPase | 6 (79) | 5.60 (2.3e-2) |

**Supplementary Table 3:**
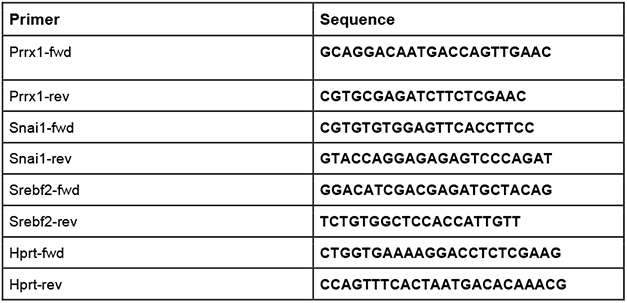
Primer sequences used for qPCR.

## References

Ashburner, M., C. A. Ball, J. A. Blake, D. Botstein, H. Butler, J. M. Cherry, A. P. Davis, et al. 2000. “Gene Ontology: Tool for the Unification of Biology. The Gene Ontology Consortium.” Nature Genetics 25 (1): 25–29.

Biddy, Brent A., Wenjun Kong, Kenji Kamimoto, Chuner Guo, Sarah E. Waye, Tao Sun, and Samantha A. Morris. 2018. “Single-Cell Mapping of Lineage and Identity in Direct Reprogramming.” Nature 564 (7735): 219.

Bray, Nicolas L., Harold Pimentel, Páll Melsted, and Lior Pachter. 2016. “Near-Optimal Probabilistic RNA-Seq Quantification.” Nature Biotechnology 34 (5): 525–27.

Briscoe, James, and Pascal P. Thérond. 2013. “The Mechanisms of Hedgehog Signalling and Its Roles in Development and Disease.” Nature Reviews. Molecular Cell Biology 14 (7): 416–29.

Butler, Andrew, Paul Hoffman, Peter Smibert, Efthymia Papalexi, and Rahul Satija. 2018. “Integrating Single-Cell Transcriptomic Data across Different Conditions, Technologies, and Species.” Nature Biotechnology 36 (5): 411–20.

Chang, Hannah H., Martin Hemberg, Mauricio Barahona, Donald E. Ingber, and Sui Huang. 2008. “Transcriptome-Wide Noise Controls Lineage Choice in Mammalian Progenitor Cells.” Nature 453 (7194): 544–47.

Colman-Lerner, Alejandro, Andrew Gordon, Eduard Serra, Tina Chin, Orna Resnekov, Drew Endy, C. Gustavo Pesce, and Roger Brent. 2005. “Regulated Cell-to-Cell Variation in a Cell-Fate Decision System.” Nature 437 (7059): 699–706.

Eling, Nils, Michael D. Morgan, and John C. Marioni. 2019. “Challenges in Measuring and Understanding Biological Noise.” Nature Reviews. Genetics 20 (9): 536–48.

Emert, Benjamin L., Christopher J. Cote, Eduardo A. Torre, Ian P. Dardani, Connie L. Jiang, Naveen Jain, Sydney M. Shaffer, and Arjun Raj. 2021. “Variability within Rare Cell States Enables Multiple Paths toward Drug Resistance.” Nature Biotechnology 39 (7): 865–76.

Francesconi, Mirko, Bruno Di Stefano, Clara Berenguer, Luisa de Andrés-Aguayo, Marcos Plana-Carmona, Maria Mendez-Lago, Amy Guillaumet-Adkins, et al. 2019. “Single Cell RNA-Seq Identifies the Origins of Heterogeneity in Efficient Cell Transdifferentiation and Reprogramming.” eLife 8 (March). https://doi.org/10.7554/eLife.41627.

Gene Ontology Consortium. 2021. “The Gene Ontology Resource: Enriching a GOld Mine.” Nucleic Acids Research 49 (D1): D325–34.

Graf, Thomas, and Tariq Enver. 2009. “Forcing Cells to Change Lineages.” Nature 462 (7273): 587–94.

Guido, Nicholas J., Philina Lee, Xiao Wang, Timothy C. Elston, and J. J. Collins. 2007. “A Pathway and Genetic Factors Contributing to Elevated Gene Expression Noise in Stationary Phase.” Biophysical Journal 93 (11): L55–57.

Hormoz, Sahand, Zakary S. Singer, James M. Linton, Yaron E. Antebi, Boris I. Shraiman, and Michael B. Elowitz. 11/2016. “Inferring Cell-State Transition Dynamics from Lineage Trees and Endpoint Single-Cell Measurements.” Cell Systems 3 (5): 419–33.e8.

Horton, J. D., I. Shimomura, M. S. Brown, R. E. Hammer, J. L. Goldstein, and H. Shimano. 1998. “Activation of Cholesterol Synthesis in Preference to Fatty Acid Synthesis in Liver and Adipose Tissue of Transgenic Mice Overproducing Sterol Regulatory Element-Binding Protein-2.” The Journal of Clinical Investigation 101 (11): 2331–39.

Huang, Pengxiang, Daniel Nedelcu, Miyako Watanabe, Cindy Jao, Youngchang Kim, Jing Liu, and Adrian Salic. 2016. “Cellular Cholesterol Directly Activates Smoothened in Hedgehog Signaling.” Cell 166 (5): 1176–87.e14.

Huang, Pengxiang, Sanduo Zheng, Bradley M. Wierbowski, Youngchang Kim, Daniel Nedelcu, Laura Aravena, Jing Liu, Andrew C. Kruse, and Adrian Salic. 2018. “Structural Basis of Smoothened Activation in Hedgehog Signaling.” Cell 174 (2): 312–24.e16.

Iwamoto, Kazunari, Yuki Shindo, and Koichi Takahashi. 2016. “Modeling Cellular Noise Underlying Heterogeneous Cell Responses in the Epidermal Growth Factor Signaling Pathway.” PLoS Computational Biology 12 (11): e1005222.

Kempe, Hermannus, Anne Schwabe, Frédéric Crémazy, Pernette J. Verschure, and Frank J. Bruggeman. 2015. “The Volumes and Transcript Counts of Single Cells Reveal Concentration Homeostasis and Capture Biological Noise.” Molecular Biology of the Cell 26 (4): 797–804.

Kinnebrew, Maia, Ellen J. Iverson, Bhaven B. Patel, Ganesh V. Pusapati, Jennifer H. Kong, Kristen A. Johnson, Giovanni Luchetti, et al. 2019. “Cholesterol Accessibility at the Ciliary Membrane Controls Hedgehog Signaling.” Edited by Duojia Pan, Marianne E. Bronner, and Stacey K. Ogden. eLife 8 (October): e50051.

Kiviet, Daniel J., Philippe Nghe, Noreen Walker, Sarah Boulineau, Vanda Sunderlikova, and Sander J. Tans. 2014. “Stochasticity of Metabolism and Growth at the Single-Cell Level.” Nature 514 (7522): 376–79.

Kong, Jennifer H., Christian Siebold, and Rajat Rohatgi. 2019. “Biochemical Mechanisms of Vertebrate Hedgehog Signaling.” Development 146 (10). https://doi.org/10.1242/dev.166892.

Kong, Wenjun, Brent A. Biddy, Kenji Kamimoto, Junedh M. Amrute, Emily G. Butka, and Samantha A. Morris. 2020. “CellTagging: Combinatorial Indexing to Simultaneously Map Lineage and Identity at Single-Cell Resolution.” Nature Protocols 15 (3): 750–72.

Larsson, Ida, Erika Dalmo, Ramy Elgendy, Mia Niklasson, Milena Doroszko, Anna Segerman, Rebecka Jörnsten, Bengt Westermark, and Sven Nelander. 2021. “Modeling Glioblastoma Heterogeneity as a Dynamic Network of Cell States.” Molecular Systems Biology 17 (9): e10105.

Lee, Raymond Teck Ho, Zhonghua Zhao, and Philip W. Ingham. 2016. “Hedgehog Signalling.” Development 143 (3): 367–72.

Li, Mingjie, Nada Husic, Ying Lin, and B. Joy Snider. 2012. “Production of Lentiviral Vectors for Transducing Cells from the Central Nervous System.” Journal of Visualized Experiments: JoVE, no. 63 (May): e4031.

Love, Michael I., Wolfgang Huber, and Simon Anders. 2014. “Moderated Estimation of Fold Change and Dispersion for RNA-Seq Data with DESeq2.” Genome Biology 15 (12): 550.

Luchetti, Giovanni, Ria Sircar, Jennifer H. Kong, Sigrid Nachtergaele, Andreas Sagner, Eamon F. X. Byrne, Douglas F. Covey, Christian Siebold, and Rajat Rohatgi. 2016. “Cholesterol Activates the G-Protein Coupled Receptor Smoothened to Promote Hedgehog Signaling.” Edited by Duojia Pan. eLife 5 (October): e20304.

McInnes, Leland, John Healy, and James Melville. 2018. “UMAP: Uniform Manifold Approximation and Projection for Dimension Reduction.” 1802.03426 [cs, Stat], February. http://arxiv.org/abs/1802.03426.

Meerbrey, Kristen L., Guang Hu, Jessica D. Kessler, Kevin Roarty, Mamie Z. Li, Justin E. Fang, Jason I. Herschkowitz, et al. 2011. “The pINDUCER Lentiviral Toolkit for Inducible RNA Interference in Vitro and in Vivo.” Proceedings of the National Academy of Sciences of the United States of America 108 (9): 3665–70.

Mi, Huaiyu, Anushya Muruganujan, Dustin Ebert, Xiaosong Huang, and Paul D. Thomas. 2019. “PANTHER Version 14: More Genomes, a New PANTHER GO-Slim and Improvements in Enrichment Analysis Tools.” Nucleic Acids Research 47 (D1): D419–26.

Miyanari, Yusuke, and Maria-Elena Torres-Padilla. 2012. “Control of Ground-State Pluripotency by Allelic Regulation of Nanog.” Nature 483 (7390): 470–73.

Neves, Ricardo Pires das, Nick S. Jones, Lorena Andreu, Rajeev Gupta, Tariq Enver, and Francisco J. Iborra. 2010. “Connecting Variability in Global Transcription Rate to Mitochondrial Variability.” PLoS Biology 8 (12): e1000560.

Padovan-Merhar, Olivia, Gautham P. Nair, Andrew G. Biaesch, Andreas Mayer, Steven Scarfone, Shawn W. Foley, Angela R. Wu, L. Stirling Churchman, Abhyudai Singh, and Arjun Raj. 2015. “Single Mammalian Cells Compensate for Differences in Cellular Volume and DNA Copy Number through Independent Global Transcriptional Mechanisms.” Molecular Cell 58 (2): 339–52.

Pliner, Hannah A., Jay Shendure, and Cole Trapnell. 2019. “Supervised Classification Enables Rapid Annotation of Cell Atlases.” Nature Methods 16 (10): 983–86.

Pusapati, Ganesh V., Jennifer H. Kong, Bhaven B. Patel, Arunkumar Krishnan, Andreas Sagner, Maia Kinnebrew, James Briscoe, L. Aravind, and Rajat Rohatgi. 2018. “CRISPR Screens Uncover Genes That Regulate Target Cell Sensitivity to the Morphogen Sonic Hedgehog.” Developmental Cell 44 (1): 113–29.e8.

Qiu, Xiaojie, Qi Mao, Ying Tang, Li Wang, Raghav Chawla, Hannah A. Pliner, and Cole Trapnell. 2017. “Reversed Graph Embedding Resolves Complex Single-Cell Trajectories.” Nature Methods 14 (10): 979–82.

Radhakrishnan, Arun, Rajat Rohatgi, and Christian Siebold. 2020. “Cholesterol Access in Cellular Membranes Controls Hedgehog Signaling.” Nature Chemical Biology 16 (12): 1303–13.

Saelens, Wouter, Robrecht Cannoodt, Helena Todorov, and Yvan Saeys. 2019. “A Comparison of Single-Cell Trajectory Inference Methods.” Nature Biotechnology 37 (5): 547–54.

Shaffer, Sydney M., Margaret C. Dunagin, Stefan R. Torborg, Eduardo A. Torre, Benjamin Emert, Clemens Krepler, Marilda Beqiri, et al. 2017. “Rare Cell Variability and Drug-Induced Reprogramming as a Mode of Cancer Drug Resistance.” Nature 546 (7658): 431–35.

Shaffer, Sydney M., Benjamin L. Emert, Raúl A. Reyes Hueros, Christopher Cote, Guillaume Harmange, Dylan L. Schaff, Ann E. Sizemore, et al. 2020. “Memory Sequencing Reveals Heritable Single-Cell Gene Expression Programs Associated with Distinct Cellular Behaviors.” Cell 182 (4): 947–59.e17.

Shaffer, Sydney M., Benjamin L. Emert, Ann E. Sizemore, Rohit Gupte, Eduardo Torre, Danielle S. Bassett, and Arjun Raj. 2018. “Memory Sequencing Reveals Heritable Single Cell Gene Expression Programs Associated with Distinct Cellular Behaviors.” Systems Biology. http://biorxiv.org/lookup/doi/10.1101/379016.

Sigal, Alex, Ron Milo, Ariel Cohen, Naama Geva-Zatorsky, Yael Klein, Yuvalal Liron, Nitzan Rosenfeld, Tamar Danon, Natalie Perzov, and Uri Alon. 11/2006. “Variability and Memory of Protein Levels in Human Cells.” Nature 444 (7119): 643–46.

Spencer, Sabrina L., Suzanne Gaudet, John G. Albeck, John M. Burke, and Peter K. Sorger. 2009. “Non-Genetic Origins of Cell-to-Cell Variability in TRAIL-Induced Apoptosis.” Nature 459 (7245): 428–32.

Street, Kelly, Davide Risso, Russell B. Fletcher, Diya Das, John Ngai, Nir Yosef, Elizabeth Purdom, and Sandrine Dudoit. 2018. “Slingshot: Cell Lineage and Pseudotime Inference for Single-Cell Transcriptomics.” BMC Genomics 19 (1): 477.

Stuart, Tim, Andrew Butler, Paul Hoffman, Christoph Hafemeister, Efthymia Papalexi, William M. Mauck 3rd, Yuhan Hao, Marlon Stoeckius, Peter Smibert, and Rahul Satija. 2019. “Comprehensive Integration of Single-Cell Data.” Cell 177 (7): 1888–1902.e21.

Stumpf, Patrick S., Rosanna C. G. Smith, Michael Lenz, Andreas Schuppert, Franz-Josef Müller, Ann Babtie, Thalia E. Chan, et al. 2017. “Stem Cell Differentiation as a Non-Markov Stochastic Process.” Cell Systems 5 (3): 268–82.e7.

Topolewski, Piotr, Karolina E. Zakrzewska, Jarosław Walczak, Karol Nienałtowski, Gerhard Müller-Newen, Abhyudai Singh, and Michał Komorowski. 2022. “Phenotypic Variability, Not Noise, Accounts for Most of the Cell-to-Cell Heterogeneity in IFN-γ and Oncostatin M Signaling Responses.” Science Signaling 15 (721): eabd9303.

Torres-Padilla, Maria-Elena, and Ian Chambers. 2014. “Transcription Factor Heterogeneity in Pluripotent Stem Cells: A Stochastic Advantage.” Development 141 (11): 2173–81.

Trapnell, Cole. 2015. “Defining Cell Types and States with Single-Cell Genomics.” Genome Research 25 (10): 1491–98.

Trapnell, Cole, Davide Cacchiarelli, Jonna Grimsby, Prapti Pokharel, Shuqiang Li, Michael Morse, Niall J. Lennon, Kenneth J. Livak, Tarjei S. Mikkelsen, and John L. Rinn. 4/2014. “The Dynamics and Regulators of Cell Fate Decisions Are Revealed by Pseudotemporal Ordering of Single Cells.” Nature Biotechnology 32 (4): 381–86.

“UniProt: The Universal Protein Knowledgebase in 2021.” 2021. Nucleic Acids Research 49 (D1): D480–89.

Zopf, C. J., Katie Quinn, Joshua Zeidman, and Narendra Maheshri. 2013. “Cell-Cycle Dependence of Transcription Dominates Noise in Gene Expression.” PLoS Computational Biology 9 (7): e1003161.

